# Identification of a novel deltavirus in Boa constrictor

**DOI:** 10.1101/429753

**Authors:** Udo Hetzel, Leonóra Szirovicza, Teemu Smura, Barbara Prähauser, Olli Vapalahti, Anja Kipar, Jussi Hepojoki

## Abstract

Human hepatitis D virus (hHDV) forms the genus *Deltavirus* which has not been assigned to any virus family. hHDV is a satellite virus and needs hepatitis B virus (HBV) to make infectious particles. Deltaviruses are thought to have evolved in humans, since they have thus far not been identified elsewhere. Herein we report, prompted by a recent observation of HDV-like agent in birds, the identification of a deltavirus in a snake (*Boa constrictor*) designated as snake-HDV (sHDV). The circular 1711 nt RNA genome of the newly identified virus resembles hHDV in its coding strategy and size. We discovered sHDV when performing a meta-transcriptomic study on brain samples of two boas with central nervous system signs. We did not identify accompanying HBV-like sequences in the samples. By sequence comparison, the putative hepatitis D antigen (HDAg) of sHDV, encoded by one of the two open reading frames (ORFs), is roughly 50% identical to the previously known HDAgs. We used antiserum raised against recombinant snake HDAg to demonstrate a broad viral target cell spectrum, ranging from neurons over epithelial cells to leukocytes. Using RT-PCR, we detected sHDV RNA also in the liver and blood of the two snakes and could show sHDV infection not only in two of their juvenile offspring, but also in a water python (*Liasis mackloti savuensis*) in the same snake colony, indicating potentially vertical and horizontal transmission. The finding of abundant sHDAg in several tissues suggests that sHDV actively replicated in the studied animals. Our findings suggest that sHDV spread may not be restricted to hepadnavirus co-infection. This would imply that deltaviruses may employ other enveloped viruses for producing infectious particles.

## Main Text

Hepatitis D virus (HDV) forms and is so far the sole member of the genus *Deltavirus* (Le Gal et al., 2006). HDV has only been found in humans, and it is represented by eight distinct genotypes (Le Gal et al., 2006; Lempp et al., 2016). In fact, HDV is hypothesized to have evolved within the human host (Littlejohn et al., 2016). HDV has a negative-sense single-stranded circular RNA genome of 1,672–1,697 nucleotides which is highly self-complementary (Lempp et al., 2016; Wang et al., 1986). The processing (autocatalytic cleavage of multimeric genomic and antigenomic RNAs and ligation of monomers) of the genome is mediated by genomic and antigenomic ribozymes (Been and Wickham, 1997; Lempp et al., 2016). HDV only encodes two proteins, the small and large hepatitis delta antigens (S- and L-HDAg), which are identical in amino acid sequence except that the L-HDAg contains 19 additional amino acid residues at its C-terminus (Tseng and Lai, 2009). The S-HDAg is needed for RNA replication and the L-HDAg is involved in virus assembly (Tseng and Lai, 2009). The virus requires hepatitis B virus (HBV) for egress and formation of infectious particles comprising a ribonucleoprotein formed of the circular RNA genome and HDAgs within an envelope decorated with HBV S antigen (Lempp et al., 2016; Rizzetto et al., 1980). HDV replicates in the nucleus and the evidence suggests that cellular DNA-dependent RNA polymerase II mediates HDV RNA replication (Tseng and Lai, 2009). Patients with chronic HBV and HDV co-infection are at great risk of developing liver cirrhosis and hepatocellular carcinoma, particularly in the case of superinfection with HDV in a chronically HBV infected patient (Lempp et al., 2016). The HDV prevalence among HBV carriers is estimated to be around 5% (Lempp et al., 2016), however, there is a great geographical and genotype-specific variation in HDV prevalence (Hughes et al., 2011). Very recently, sequence data showing presence of a divergent HDV-like agent was reported in ducks, without any traces of duck orthohepadnavirus (Wille et al., 2018). This prompted us to report our findings of a HDV-like agent which we discovered earlier this year.

The animals included in this study were submitted to the Institute of Veterinary Pathology, Vetsuisse Faculty, University of Zurich, Switzerland for a diagnostic post mortem examination upon the owner’s request. We applied the Animals Scientific Procedures Act 1986 (ASPA), schedule 1 (http://www.legislation.gov.uk/ukpga/1986/14/schedule/1) procedure to euthanize the snakes. Euthanasia and diagnosis-motivated necropsies are both routine veterinary procedures, and thus ethical permissions were not required.

The animals carrying the snake HDV (sHDV) were a *Boa constrictor sabogae* breeding pair with their joint offspring (F2, F3) and a waterpython (*Liasis mackloti savuensis*) from the same colony (Table 1). The parental animals (nos. 1 and 3) had originally been imported from Panama to Italy, from where they were sold to a private owner in Switzerland. All snakes had shown mild neurological signs, which were suspected to be associated with boid inclusion body disease (BIBD). Confirmation of BIBD was achieved by examination of blood smears. After euthanasia, the diagnoses was confirmed by histological examination of formalin-fixed paraffin-embedded samples of brain and other tissues. Apart from the waterpython which suffered from a chronic hepatitis, none of the snakes exhibited other histopathological changes. We used RNA extracted from a freshly frozen brain sample of the parental animals for next-generation sequencing (NGS) library preparation as described in (Keller et al., 2017). The sequencing by Illumina MiSeq platform with MiSeq Reagent Kit v3 (Illumina) 2x300 cycles yielded 825,933 paired end reads, and removal of reads matching to snakes genome (*Python bivitattus*) reduced the number of paired end reads to 401,141. We performed *de novo* assembly using MIRA version 4.9.5 (http://mira-assembler.sourceforge.net/) on CSC (IT Center for Science Ltd., Finland) Taito supercluster. One of the contigs with high coverage (130,902 reads in total, corresponding to 7.92% of all reads) appeared to be circular. The contig contained three repeats of 1,711 nt sequence (GenBank, accession no. MH988742) with two open reading frames (ORFs), one in sense and the other in anti-sense orientation. The genome of the newly identified sHDV and the sequencing coverage are shown in Fig. 1A. By BLAST analysis (https://blast.ncbi.nlm.nih.gov/Blast.cgi) the other ORF of 199 amino acids was identified as snake HDAg (sHDAg) with 50% amino acid identity to S-HDAg (GenBank accession no. AGI51675.1). The ORF2 contains a stretch resembling the DUF3343 (domain of unknown function) by HMMER3 search in SMART (Simple Modular Architecture Research Tool available at http://smart.embl-heidelberg.de/), but with no apparent other homologies to known proteins. The secondary structure of the genome generated using RNAstructure webserver (Bellaousov et al., 2013) shows 73% self-complementarity, which is close to the 74% reported for known hHDVs (Lempp et al., 2016). By GC content (53.3%) the sHDV lies between the newly reported avian HDV-like sequence (51%) (Wille et al., 2018) and human HDV (hHDV, 60%) (Wang et al., 1986).

**Table 1.** Animals included in the study.

| **No.** | **Animal** | **Additional information** | **Delta-PCR (organs tested)** | **Delta-IH (organs positive)** |
| --- | --- | --- | --- | --- |
| 1 | *B. c. sabogae*, female, >12 y | Maternal animal; NGS | **Pos** (liver, blood, brain) | **Pos** (brain, liver, lung, spleen, kidney) |
| 2 | *B. c. sabogae*, male, 2 y | Offspring of #1 and #3 (F3) | **Neg** (liver) | **Neg** |
| 3 | *B. c. sabogae*, male, > 12y | Paternal animal; NGS | **Pos** (liver) | **Pos** (brain, liver, lung, spleen, kidney) |
| 4 | *Liasis mackloti savuensis*, male, > 10 y | Housed in the same room as #1 and #3; chronic hepatitis, no BIBD | **Pos** (liver) | **Pos** (liver) |
| 5 | *B. c. sabogae*, female, 3 y | Offspring of #1 and #2 (F2) | **Pos** (liver) | **Pos** (brain, liver, spleen, kidney) |
| 6 | *B. c. sabogae*, male, 3 y | Offspring of #1 and #2 (F2) | **Pos** (liver) | **Pos** (spleen, liver) |
| 7 | *B. c. constrictor*, female, ca. 6 y | Housed in the same room as #1 and #3; no BIBD | **Neg** (liver) | **Neg** |
| 8 | *B. c. sabogae*, male, 2 y | Offspring of #1 and #3 (F3) | **Neg** (liver) | **Neg** |
| 9 | *Sanzinia madagascariensis*, female, 15 y | Control animal; from different breeder; cutaneous mycosis, no BIBD | **Neg** (liver) | **Neg** |
| 10 | *B. c. sabogae*, female, 3 y | Offspring of #1 and #3 (F2) | **Pos** (liver) | **Pos** (liver, brain) |
| 11 | *B. c. sabogae*, male, 3 y | Offspring of #1 and #3 (F2) | **Pos** (liver) | **Pos** (spleen, liver, brain) |

B.c. - Boa constrictor, y - years, NGS - next generation sequencing, F2/F3 - second/third clutch of breeding pair (#1 and #3)

**Figure 1.**
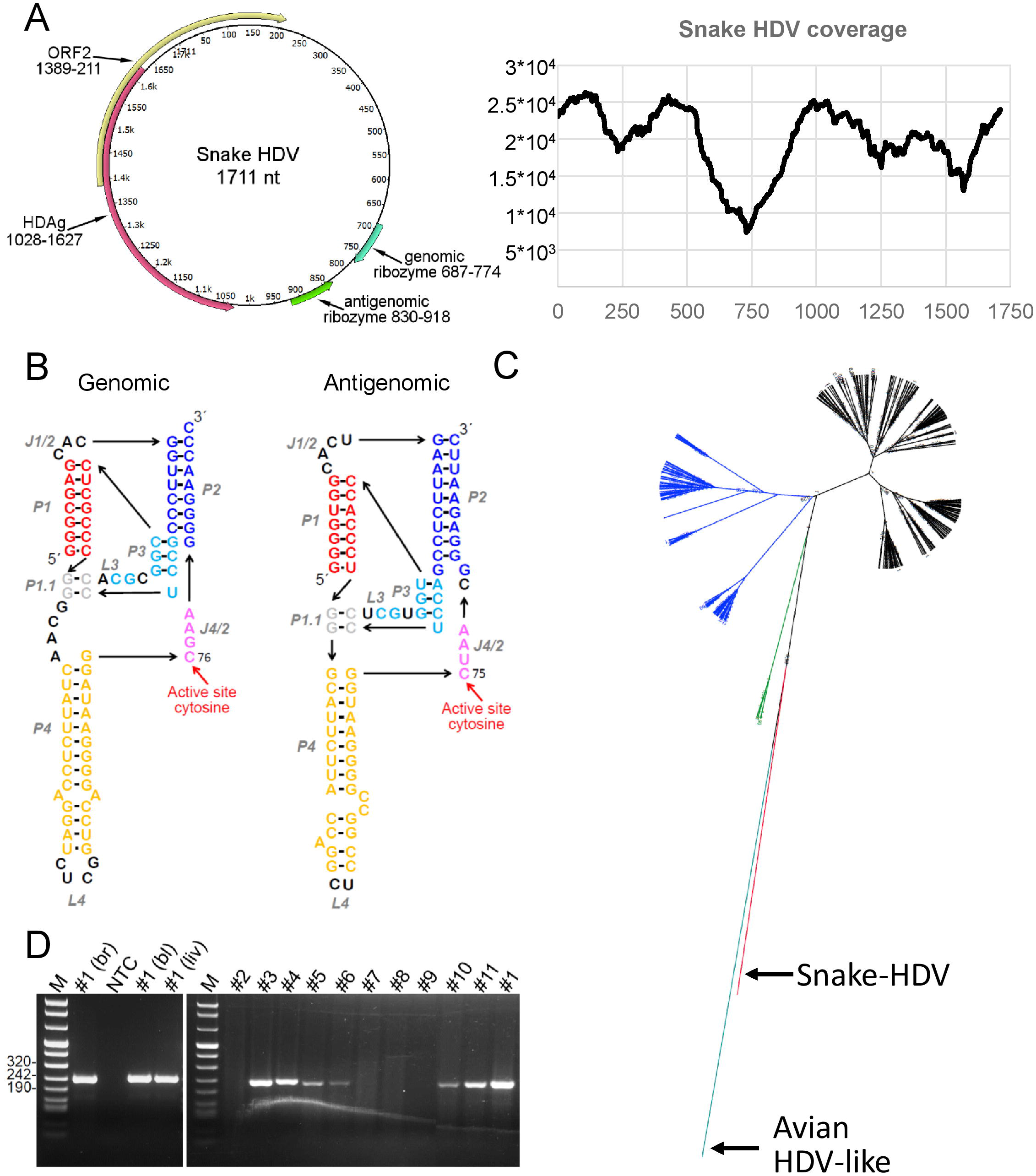
Genome organization, sequencing coverage, schematic ribozyme structure and phylogenetic analysis of snake HDV. **A)** Schematic presentation of circular RNA genome and sequencing coverage for snake HDV. The genome shows two open reading frames (ORFs), the ORF in antigenomic orientation spanning nucleotide residues 1028–1627 encodes a 199 amino acid protein which by BLAST analysis represents the HDAg. The ORF2 is in genomic orientation and spans residues 1389–211 encodes 177 amino acid protein, which by BLAST analysis did not yield significant hits (35% identity over 66 amino acids, E-value 5, to ferritin-like protein from *Candidatus Nitrososphaera evergladensis* SR1, AIF82718.1). SMART (Simple Modular Architecture Research Tool available at http://smart.embl-heidelberg.de/) analysis showed the putative protein to have 2 transmembrane helices, and a DUF3343 (domain of unknown function) domain with E-value 0.013. The genomic and antigenomic ribozymes identified by sequence alignments to known HDVs are respectively located at 687–744 and 830–918. The graph shows sequencing coverage (on Y-axis) in respect to each nucleotide position (on X-axis) of snake HDV from the original brain sample, the coverage ranges from 7368 (at nt position 729) to 26304 fold. **B)** Models for the secondary structures of the genomic and antigenomic ribozymes identified in snake HDV, the presentation format is adopted from review by Webb and Luptak (Webb and Luptak, 2011) which was also used by Wille et al. (Wille et al., 2018). The abbreviations are: P = paired region, J = joining region, L = loop. Both genomic and antigenomic ribozymes are structurally close to their human HDV counter parts described in (Webb and Luptak, 2011) and they are identical at the following regions: active site, P1.1 and P3. The cleavage by the ribozyme occurs at the 5′ end. **C)** Phylogenetic analysis of human, avian, and reptile HDAgs. The phylogenetic analysis was done using Bayesian MCMC method implemented in MrBayes 3.1.2 (Ronquist et al., 2012) with JTT model of substitution with gamma distributed rate variation among sites. HDV genotype 1 is in black, HDV genotype 2 in blue, HDV genotype 3 in green, avian HDV-like sequence in cyan, and snake-HDV in red. **D)** RT-PCR results of snake tissues. The gel on left shows RT-PCR products obtained for snake 1 (table 1) from different tissues: br = brain, bl = blood, liv = liver. NTC = non template control, M is DNA ladder. The gel on right shows RT-PCR products obtained from liver sample, the animal numbering is according to table 1, animal #1 serves as a positive control.

By aligning the nucleotide sequence of sHDV with hHDVs, we were able to locate the genomic and antigenomic ribozymes (Fig. 1B). The ribozymes share several features with the hHDV counterparts including the active site and surrounding nt residues. The phylogenetic analysis of amino acid sequences of hHDAgs shows that the sHDV and avian HDV-like agents (Wille et al., 2018) are divergent from the hHDV. sHDAg forms a sister clade to hHDAgs, whereas HDAg of the avian HDV-like agent forms an outgroup for these (Fig. 1C).

RT-PCR targeting the nt region 1139–1374 was set up using the 5′-GGATTGTCCCTCCAGAGGGTTC-3′ (fwd) and 5′-GCTCGAGGCTACCACCGAAAG-3′ (rev) primer pair. We performed conventional RT-PCR as described in (Keller et al., 2017) using RNA extracted from freshly frozen liver samples as the template as described in (Dervas et al., 2017). We found the parental animals and four of their seven offspring as well as the waterpython (no. 4) to be sHDV-infected (Table 1 and Fig. 1D). The latter animal had been housed in the same room as the boa breeding pair for several years, similar to an adult *B. constrictor constrictor* (no. 7) that was tested negative for sHDV. A Madagascar tree boa (*Sanzinia madagascariensis*) without BIBD from a different breeder was equally negative (Table 1).

To produce an antibody against the sHDAg, we used Champion pET101 Directional TOPO Expression Kit (Thermo Scientific) to clone and express the recombinant sHDAg with C-terminal hexa-histidine tag. We designed primers (5′-CACCATGGAAACTCCATCCAAGAAGC-3′[fwd] and 5′-CGGGAACATTTTGTCACCCCTCAC-3′ [rev]) according to the manufacturer’s instructions to PCR amplify sHDAg ORF from the brain sample used for NGS library preparation. We did the protein expression similarly as described in (Dervas et al., 2017), but performed the purification under native conditions using Ni-NTA agarose (QIAGEN) according to manufacturer’s instructions. Rabbit antiserum against the recombinant protein was prepared by BioGenes GmbH as described in (Dervas et al., 2017). Immunohistology then served to detect sHDAg expression in the formalin-fixed and paraffin-embedded tissues (brain, liver, lung, kidney, spleen) of the examined snakes, using the anti-sHDAg antiserum (see above). We used the EnVision HRP detection system (Dako) as described (Dervas et al., 2017), citrate buffer (pH 6.0 at 98°C, 10 min) for antigen retrieval of the sections, and anti-sHDAg serum at 1:5,000 dilution in Dako dilution buffer. Consecutive sections incubated with the pre-immune serum instead of the specific primary antibody served as negative controls.

In both parental boas, we found sHDAg to be intensely expressed within the cell body and processes of numerous neurons in all brain regions (Fig. 2A), in individual hepatocytes in the liver (Fig. 2B), in a proportion of tubular epithelial cells in the kidney (Fig. 2C), in occasional epithelial cells in the lung (Fig. 2D), and in leukocytes (mainly consistent with macrophages) in the spleen (Fig. 2E). All tissues also showed evidence of viral antigen expression in occasional vascular endothelial cells and some leukocytes (Fig. 2). These findings suggest active replication. Of the seven juvenile offspring tested, we found the four RT-PCR-positive animals to also be positive by immunohistology, though mainly with a more limited expression (Table 1). The RT-PCR positive waterpython exhibited patchy sHDAg expression in the liver. The three RT-PCR negative boa offspring as well as the RT-PCR negative control animal were also negative in the immunohistology.

**Figure 2.**
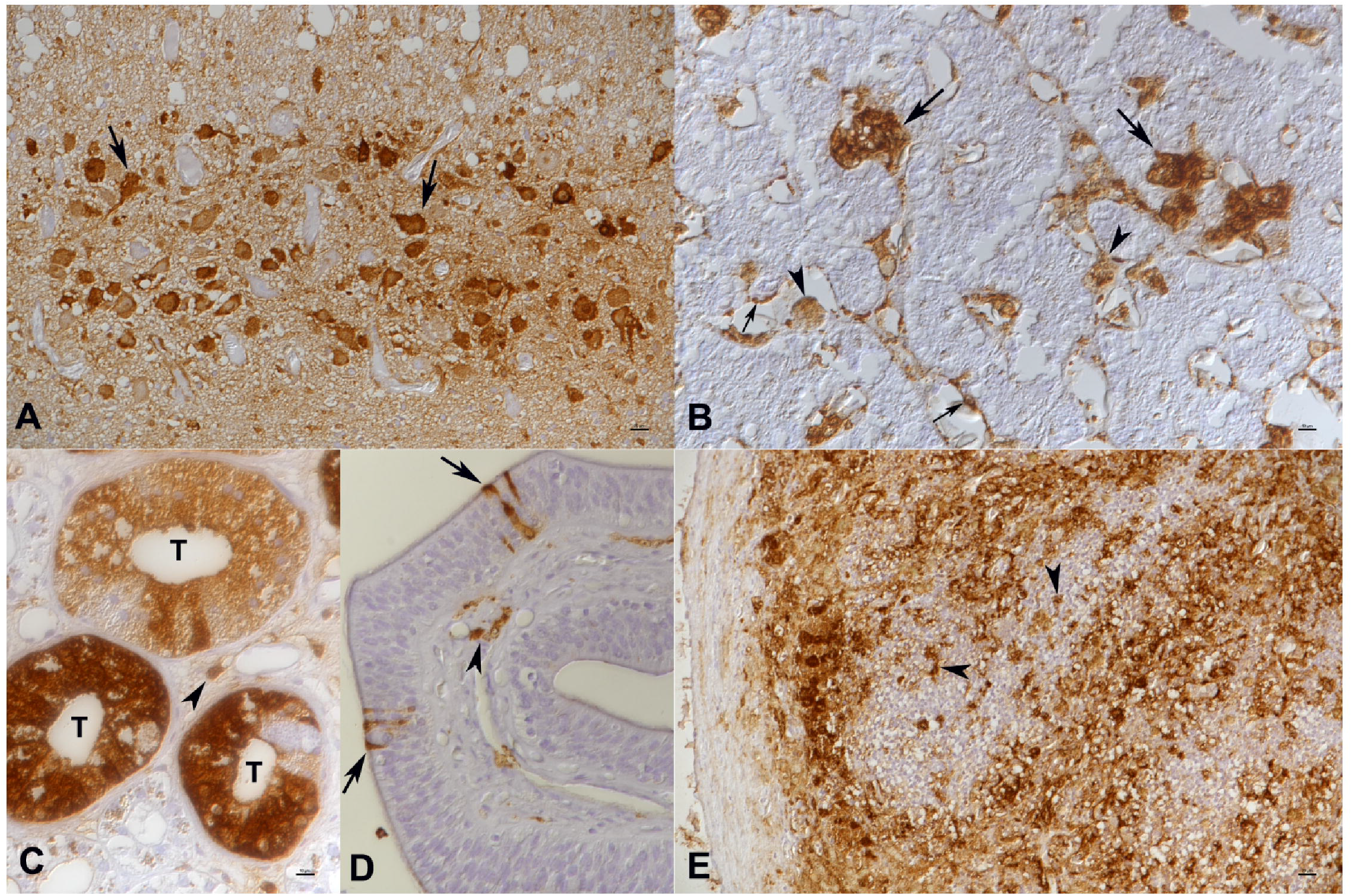
*Boa constrictor*, female adult. **A)** Brain. Viral antigen is expressed in nucleus, cytoplasm and cell processes of numerous neurons. **B)** Liver. Individual hepatocytes (large arrows) are strongly positive, and macrophages (arrowheads) and endothelial cells (small arrows) are found to also express viral antigen. **C)** Kidney. In a group of tubules (T), the majority of epithelial cells exhibit variably intense viral antigen expression. Occasional leukocytes in the interstitium (arrowhead) are also positive. **D)** Lung. There are several individual positive epithelial cells (arrows); some subepithelial leukocytes are also found to express viral antigen (arrowhead). **E)** Spleen. There is extensive viral antigen expression. Positive cells often have the morphology of macrophages (arrowheads). Immunohistology for sHDAg. Horseradish peroxidase method, haematoxylin counterstain.

Herein we provide the first evidence of actively replicating deltavirus in species other than man. Together with the recent report by Wille et. al. (Wille et al., 2018) our study also suggests that deltaviruses are in fact likely found across several taxa. Our histological examination shows that the tropism of sHDV is broad and not limited to liver and blood. In fact, the detection of sHDAg in the renal tubular epithelium and lung epithelial cells indicates that the virus can be shed with secretions. We could not associate the infection with cytopathic changes, but need to undertake further studies to assess the sHDV-related pathogenesis. The fact that we, alike Wille et al. (Wille et al., 2018), could not detect accompanying hepadnavirus challenges the current understanding of strict hepadnavirus-deltavirus association. It would seem plausible that the newly found deltaviruses use arenavirus (in case of snakes) and influenza virus (in case of birds) co-infection to obtain the lipid envelope to make infectious particles. These findings open up a multitude of avenues in deltavirus research.

## Acknowledgements

The authors are grateful to Sabina Wunderlin and Barbara Prähauser, Histology Laboratory, Institute of Veterinary Pathology, Vetsuisse Faculty, University of Zurich, for excellent technical support. This work was supported by Academy of Finland grants to JH (grant numbers 130861 and 1314119).

